# Divergent and dominant anti-inflammatory effects of patient-derived anti-citrullinated protein antibodies (ACPA) in arthritis development

**DOI:** 10.1101/2022.09.30.510377

**Authors:** Bruno Raposo, Marcelo Afonso, Lena Israelsson, Heidi Wähämaa, Ragnhild Stålesen, Fredrik Wermeling, Aase Hensvold, Caroline Grönwall, Bence Rethi, Lars Klareskog, Vivianne Malmström

## Abstract

About two thirds of rheumatoid arthritis (RA) patients develop anti-citrullinated protein antibodies (ACPA), which are a characteristic of the disease and sparsely observed in other clinical settings or in the healthy population. ACPA are used as a diagnostic criterion and often develop prior to diagnosis. Therefore, ACPA titers are important in the identification of individuals at risk of developing RA. Interestingly, the titers and target cross-reactivity of ACPA increase by time of diagnosis, suggesting a causality between anti-citrulline reactivity and the development of RA. However, only 50% of ACPA-positive at-risk individuals will progress to a clinical RA diagnosis. This observation suggests that there are different types of ACPA and that their relationship with RA development is more complex than presently assumed.

To address the biological effect of ACPA in the establishment and development of inflammatory arthritis, we made use of the collagen antibody-induced arthritis (CAIA) model of passive arthritis. Using seven unique patient derived monoclonal ACPA, we observed that ACPA are predominantly anti-inflammatory, with some clones (C03 and BVCA1) completely inhibiting disease development. We have also identified a clone (C04) as having a disease-prone effect, by increasing and sustaining disease prevalence. This clone had previously been reported as being pro-inflammatory in a different model of joint inflammation. The anti-inflammatory effects of C03 were dependent on FcγR, since neither F(ab’)2 or a mutated Fc-null GRLR-C03 clone could mediate disease protection. Most importantly, mice receiving polyclonal ACPA enriched from RA sera were also partially protected from disease. Looking at the cell and tissue of origin of the ACPA clones, their V(D)J sequences, Fc glycosylation pattern and fine-specificities, we could not identify a common feature between anti-inflammatory ACPA distinguishing them from those without the properties. Together, our data demonstrates that circulating ACPA in RA patients are predominantly anti-inflammatory, emphasizing the need to study ACPA repertoires in order to determine their influence on the clinical outcome of the patient or at-risk individual.

## Letter

The presence of anti-citrullinated protein antibodies (ACPA) is currently used in RA diagnosis and distinguishes the major subsets of patients. The demonstration that ACPA occur before onset of RA and associate with severe disease [1] has been used to advocate a causal role of ACPA in disease development. However, not all persistent ACPA-positive individuals progress to clinical RA, suggesting a complex relationship between ACPA and arthritis development, where ACPA displaying different inflammatory aptitudes may exist[2]. In recent years, we and others have isolated single B cells from RA patients and re-expressed monoclonal ACPAs[3,4]. These monoclonal ACPA display different properties regarding immune-mediated processes in vitro, as well as in vivo phenotypes such as pain and bone erosion[5,6]. Using the collagen antibody-induced arthritis (CAIA) model of passive arthritis, we assessed the properties of different monoclonal ACPA in vivo concerning their ability to modify the arthritic process (figure 1A). To address this question in an unbiased manner, we tested seven monoclonal ACPA expressed as murine chimeric IgG2a and displaying unique profiles of citrulline-directed fine-specificities (figure 1B and supplementary figure 1). In line with previous evidence, ACPA per se did not induce arthritis (supplementary figure 2A). Interestingly however, we observed several monoclonal ACPA inhibiting (clones mC03 and mBVCA1) or ameliorating arthritis (mB09 and mA01; figure 1B, 1C and 1D), whereas other clones showed no effect in the model (mX1604, m17D08; figure 1C and 1D), while one clone provided a slightly enhanced arthritis prevalence (mC04; figure 1C). Using a different model of joint inflammation, the mC04 monoclonal ACPA was previously shown to have an arthritis-accelerating effect[7]. Strikingly, mice receiving the mC03 clone at the peak of disease recovered almost completely from joint inflammation 48h post ACPA transfer (figure 1E). Similar results were observed with mB09, although less pronounced (supplementary figure 2B). When combining mC03 and mC04 ACPA, the anti-inflammatory effect of mC03 ACPA prevailed, i.e. no arthritis developed (supplementary figure 2C). The inhibitory effect on arthritis was not linked to any of the known ACPA clones’ fine-specificities, nor to a previously reported histone epitope associated with this effect (supplementary figure 1E)[2]. The antiinflammatory effects induced by mC03 were clearly FcγR-dependent, with both its F(ab’)2 fragments and FcγR null (GRLR-mutated) variants incapable of suppressing arthritis development (figure 1F and 1G; comparison between murine and human C03 ACPA in supplementary figure 2D). However, no parallel differences in terms of expression of activating or inhibitory FcγR in blood immune cells throughout the course of disease could be observed (supplementary figure 3). These effects also seemed independent of the ACPA Fc-glycosylation patterns, and the capacity of ACPA in activating the classical complement pathway (supplementary figure 4). Importantly, using IgG ACPA purified by affinity chromatography from a pool of sera from RA patients, we demonstrate that these polyclonal ACPA seem to be dominantly anti-inflammatory in the CAIA model, contrasting to the respective non-ACPA fraction of the sera (figure 1H).

**Figure 1.**
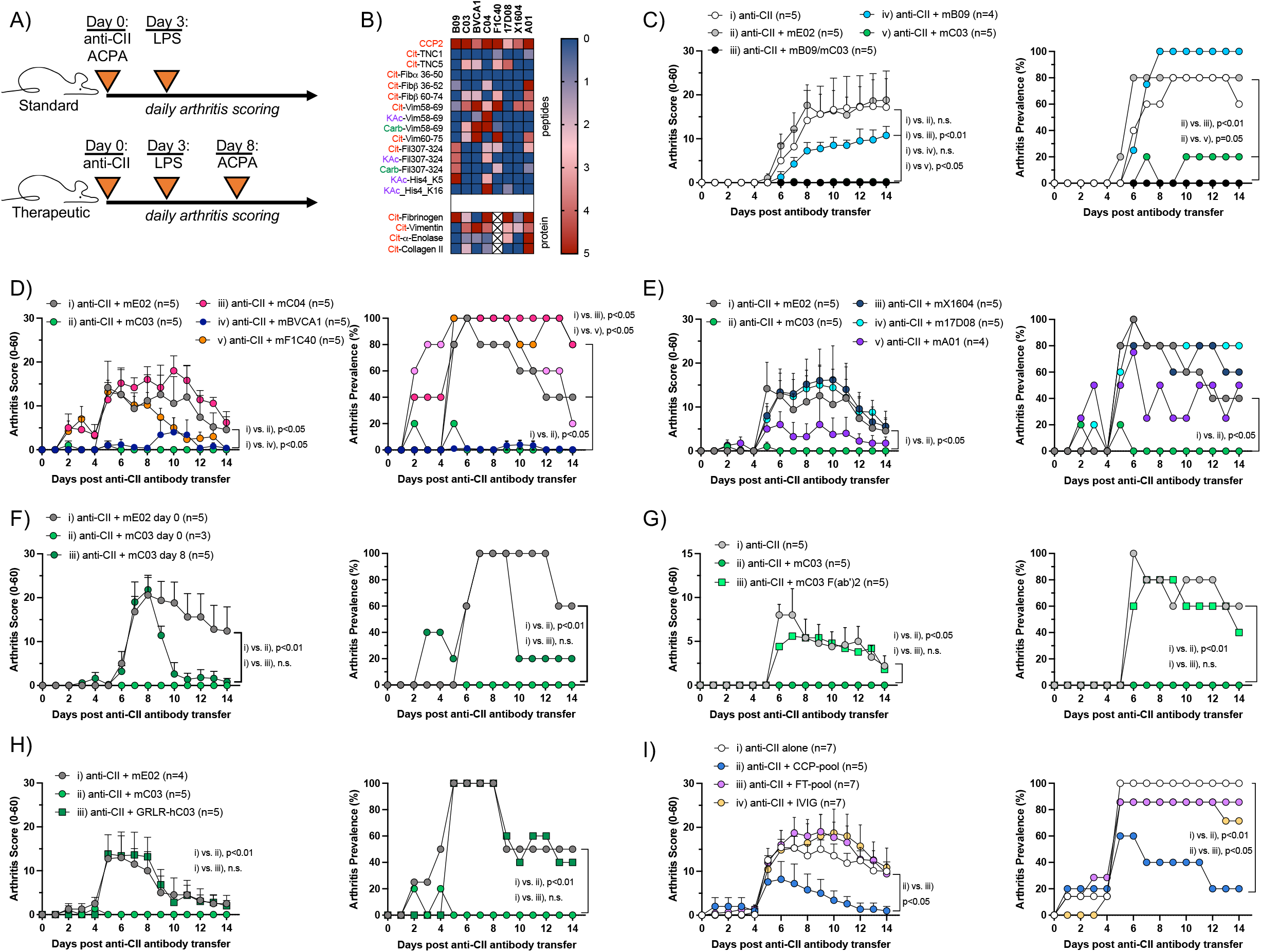
Dominant anti-inflammatory properties evidenced by distinct ACPA clones in the development of inflammatory arthritis. (A) Schematic representation of the animal model used. (B) Anti-citrullinated peptide and protein reactivity by monoclonal ACPAs used in the study. (C-I) CAIA was induced in mice by i.v. transfer of arthritogenic anti-CII antibody cocktail. Monoclonal ACPAs (C, D and E), C03 F(ab’)2 fragments (G), mutated FcγR-null GRLR-C03 ACPA (H) or polyclonal CCP-pool and respective flow-through (I) were transferred simultaneously at time of disease induction. Therapeutic effect of ACPAs was assessed by transfer of C03 ACPA at the peak of disease, day 8 (F). Boosting and synchronization of disease symptoms was done by i.p. administration of LPS 3 days post disease induction. Statistical analysis calculated by non-parametric repeated-measures Friedman test with Dunn’s multiple comparison test. Disease curves from mC03 and mE02 reference groups are identical in panels C and D due to splitting of the data into two panels for better visualization. Cit: citrullination; KAc: acetylated lysine; Carb: carbamylation; TNC: tenascin C; Fib: fibrinogen; Vim: vimentin; Fil: fillagrin; His4: histone 4; mC03: murine IgG2a C03; hC03: human IgG1 C03; FT: flow-through; IVIG: intravenous IgG preparation.

The observation that certain ACPA display a beneficial phenotype in inflammatory arthritis needs to be further understood and explored. Whereas some of the here used monoclonal ACPA have previously shown to induce symptoms such as arthralgia, bone loss or tenosynovitis that often precede onset of RA, the present demonstration of a dominant anti-inflammatory effect by certain mono- and polyclonal ACPA calls for a re-evaluation of the proposed role of these antibodies in RA. This re-evaluation must consider the heterogeneity of effects mediated by monoclonal ACPA, which together with additional immune stimuli may influence whether an ACPA-positive individual progresses to clinical RA. We acknowledge that the complete molecular mechanism mediating the ACPA anti-inflammatory effects is currently unknown to us and admittedly CAIA is not a citrullination-dependent arthritis model, but the striking effects here shown warrant further investigations. The availability of the monoclonal ACPA used here will enable such urgent investigations to take place.

---

**Supplementary figure 1 - Citrullinated antigen reactive fine-specificities by monoclonal ACPA do not associate with observed anti-inflammatory effects.** Reactivity to citrullinated and native full protein fibrinogen (A), vimentin (B), alpha-enolase (C) and type II collagen (D) by monoclonal ACPAs and respective associations with observed anti-inflammatory effects in CAIA. (E) Reactivity of tested ACPA clones against the citrulline 3 peptide from histone 4 (His4) identified by Chirivi and colleagues [*R. Chirivi et al (2021); Therapeutic ACPA inhibits NET formation: a potential therapy for neutrophil-mediated inflammatory diseases; Cellular and Molecular Immunology 18:1528]*. Nat-Fib: native fibrinogen; cit-Fib: citrullinated fibrinogen; nat-Vim: native vimentin; cit-Vim: citrullinated vimentin; nat-ENO1: native alpha-enolase; cit-ENO1: citrullinated alpha-enolase; nat-CII: native type II collagen; cit-CII: citrullinated type II collagen.

**Supplementary figure 2 – Supporting observations on how ACPAs affect inflammatory arthritis in vivo.** (A) Transfer of anti-CII antibody cocktail or a combination of two monoclonal ACPAs i.v. day 0, followed by LPS administration i.p. day 4. (B-D) CAIA was induced in mice by i.v. transfer of arthritogenic anti-CII antibody cocktail. Boosting and synchronization of disease symptoms was done by i.p. administration of LPS 3 days post disease induction. (B) mB09 ACPA transferred at day 0 or 7 to assess therapeutic potential. (C) Co-administration of anti-inflammatory mC03 and disease-prone mC04 ACPAs at day 0, or mC03 day 0 followed by mC04 day 7. (D) Comparison between C03 ACPA expressing a murine IgG2a or human IgG1 constant domain. Statistical analysis calculated by nonparametric repeated-measures Friedman test with Dunn’s multiple comparison test.

**Supplementary figure 3 – Frequency of circulating immune cells and their expression of FcγR during CAIA is not affected by administration of ACPA.** CAIA was induced in BALB/c mice followed by LPS administration day 3. mE02 or mC03 ACPA were transferred at day 0. Blood was collected at days 0, 3, 6, 9 and 13 for evaluation of immune cell frequencies in the blood, as well as their corresponding expression of FcγR1, FcγR2b, FcγR3, FcγR4. (A) Gating strategy. (B) Representation of FcγR expression in mast cells (FcεR+ c-kit+), (C) monocytes (Ly6C+Ly6C–),(D) inflammatory monocytes/macrophages (Ly6Ghi F4/80+), (E) neutrophils (Ly6G+Ly6C+), (F) DCs (CD11c+ CD19–) and (G) B cells (CD19+ CD11c–). No differences in mean fluorescence intensity (MFI) of any of the FcγR were detected between mice receiving mE02 or mC03.

**Supplementary figure 4 – Glycosylation forms and complement deposition by ACPA clones do not clearly explain observed anti-inflammatory effects.** (A) Anti-citrulline reactivity was determined through a CCP-ELISA in serum from naïve mice, mice that were submitted to CAIA (day 15 post disease induction), as well as the anti-CII arthritogenic antibody cocktail used to induce the arthritis model. (B) Complement C1q deposition induced by different ACPA clones. (C) Representation of the normalized C1q binding shown in (B) at a concentration of 1.25μg/ml monoclonal ACPA. (D) Schematic representation of major Fc glycoforms. FA2G2S1 and FA2G2S2 glycoforms were not found in any of the ACPA clones. (E) Relative abundance of each glycoform found in ACPA clones used in vivo.

## Acknowledgements

We would like to thank Federica Sallusto and Luca Piccoli for their contribution in the generation of the BVCA1 ACPA clone, and Daniel Mueller (University of Minnesota Medical School) for collaboration when generating the C04 and F1C40 ACPA clones. For glycoproteomics analysis we thank Ekaterina Mirgorodskaya at the Proteomics Core Facility at Sahlgrenska Academy, University of Gothenburg, Sweden supported by the Swedish National Infrastructure (BioMS) and SciLifeLab..

## References

1 Hair MJH, Sande MGH, Ramwadhdoebe TH, et al. Features of the Synovium of Individuals at Risk of Developing Rheumatoid Arthritis: Implications for Understanding Preclinical Rheumatoid Arthritis. Arthritis Rheumatol 2014;66:513–22. doi:10.1002/art.38273

2 Chirivi RGS, Rosmalen JWG van, Linden M van der, et al. Therapeutic ACPA inhibits NET formation: a potential therapy for neutrophil-mediated inflammatory diseases. Cell Mol Immunol 2021;18:1528–44. doi:10.1038/s41423-020-0381-3

3 Sahlström P, Hansson M, Steen J, et al. Different Hierarchies of Anti–Modified Protein Autoantibody Reactivities in Rheumatoid Arthritis. Arthritis Rheumatol 2020;72:1643–57. doi:10.1002/art.41385

4 Kissel T, Reijm S, Slot L, et al. Antibodies and B cells recognising citrullinated proteins display a broad cross-reactivity towards other post-translational modifications. Ann Rheum Dis 2020;79:472. doi:10.1136/annrheumdis-2019-216499

5 Jurczak A, Delay L, Barbier J, et al. Antibody-induced pain-like behavior and bone erosion: links to subclinical inflammation, osteoclast activity, and acid-sensing ion channel 3– dependent sensitization. Pain 2022;163:1542–59. doi:10.1097/j.pain.0000000000002543

6 Krishnamurthy A, Circiumaru A, Sun J, et al. Combination of two monoclonal ACPAs induced tenosynovitis, pain and bone loss in mice in a Peptidyl Arginine Deiminase-4 dependent manner. Arthritis Rheumatol Published Online First: 2022. doi:10.1002/art.42320

7 Titcombe PJ, Wigerblad G, Sippl N, et al. Pathogenic Citrulline-Multispecific B Cell Receptor Clades in Rheumatoid Arthritis. Arthritis Rheumatol 2018;70:1933–45. doi:10.1002/art.40590

