## Supplementary material for "Divergent and dominant anti-inflammatory effects of patient-derived anti-citrullinated protein antibodies (ACPA) in arthritis development": Material and Methods

*Origin and re-expression of patient-derived monoclonal antibodies*

Single B cells from RA patient were isolated using different methodologies and the immunoglobulin variable regions were amplified by reverse transcription and multiplex PCR and sequenced as previously described[1]. For in vitro binding studies the antibodies were subcloned as human IgG1 with human γ1, κ/λ constant regions by an off-site gene synthesis service (IDT). For in vivo studies the mAbs were subcloned as murine chimeric IgG2a constructs with murine γ2a/c, κ/λ constant regions. Monoclonal antibodies were recombinantly expressed in Expi293F cells (ThermoFisher Scientific), purified using protein G Sepharose (Cytiva), and extensively quality controlled using SDS-PAGE, ELISA, size exclusion chromatography, and endotoxin testing[1]. All clones were initially identified as ACPAs by screening using antigen microarray binding to citrullinated peptides and CCP2 by CCPlus ELISA (Svar Life Science) at 5 μg/ml. No clones had any reactivity to native peptides or unspecific polyreactivity by soluble membrane protein (SMP) assay[2]. All ACPA clones were derived from CCP2 positive RA patients. The clones 1325:01B09 (hereafter B09) and 1325:04C03 (hereafter C03) were derived from synovial plasma cells from the same ACPA+ RA patient[3]. The cells were isolated based on antibody secretion based using the FOCI method. The BVCA1 clone is derived from a blood memory B cell using and immortalization protocol[4]. The clones 37CEPT2C04 (hereafter C04) and 37CEPF1C40 (hereafter F1C40) were derived from memory blood B cells from the same ACPA+ RA patient using tetramer antigen staining and flow cytometry sorting for reactivity to citrullinated alpha enolase (CEP1 peptide)[5]. The clone L204:01A01 (hereafter A01) was derived from a lung plasma blast from an early untreated RA patient by flow cytometry sorting BAL B cells. 254:17D08 (hereafter 17D08) and 254:C7X1604 (hereafter X1604) were derived from bone marrow from the same RA patient using either flow cytometry sorting of CD138+ cells or 10X Genomic single cells transcriptomics.

Engineered human FcγR null IgG1 construct with G236R/L328R mutations (GRLR) which has previously shown not to interact with any FcγR[6], was generated by gene synthesis (IDT) and megaprimer PCR of the γ1 vector[7]. Unaffected citrulline reactivity was confirmed by CCP2 and cit-peptide ELISA.

Supplemental Table 1. Human monoclonal ACPA

| **Clone** | **Patient** | **B cell** | **Method of isolation** | **CCP2 reactivity#** | **Fab glycosylation sites** |
|---|---|---|---|---|---|
| 1325:04C03 | Established ACPA+ RA (patient A) | Synovial plasma cell | FOCI[3] | 1500 AU/ml | Yes – 2 VH; 0 VL |
| 1325:01B09 | Established ACPA+ RA (patient A) | Synovial plasma cell | FOCI[3] | 1170 AU/ml | Yes – 1 VH; 1 VL |
| BVCA1 | Established ACPA+ RA (patient B) | Blood memory | Immortalization[4] | 795 AU/ml | Yes – 0 VH; 1 VL |
| 37CEPT2C04 | Established ACPA+ RA (patient C) | CEP1+ blood memory | Antigen-specific flow cytometry sorting[5] | 3004 AU/ml | Yes – 2 VH; 1 VL |
| 37CEPF1C40 | Established ACPA+ RA (patient C) | CEP1+ blood memory | Antigen-specific flow cytometry sorting[5] | 1884 AU/ml | No |
| L204:01A01 | Early untreated RA (patient D) | Lung plasmablast | Flow cytometry sorting CD19+ cells BAL fluid | 3200 AU/ml | Yes – 1 VH; 1 VL |
| 254:17D08 | Established ACPA+ RA (patient E)* | Bone marrow plasma cell | Flow cytometry sorting CD19+ CD138+ | 432 AU/ml | Yes – 3 VH; 1 VL |
| 254:C7X1604 | Established ACPA+ RA (patient E)* | Bone marrow plasma cell | Single cells transcriptomics | 566 AU/ml | Yes – 1 VH; 2 VL |
| 1362:01E02 | ACPA- RA (patient F) | Synovial memory B cells | Flow cytometry sorting CD19+ CD27+ | 0 AU/ml | No |

BAL: bronchoalveolar lavage
* patient undergoing hip arthroplasty
### CCPlus (Svar Life Science) at 5 μg/ml hIgG1

*Collagen antibody-induced arthritis (CAIA) and antibody transfers*

Unless described otherwise, all CAIA experiments were conducted in 7 weeks old male BALB/c mice. Mice were acquired from The Jackson Laboratory (USA) or Charles River (France) and maintained in a SPF animal facility with access to water and rodent chow *ad libitum*, in environment-enriched cages. Induction of CAIA was done by i.v. transfer of 1.5 mg per mouse of a 5 monoclonal anti-type II collagen (CII) antibody cocktail (Chondrex, USA). 3 days later, mice were challenged i.p. with 25 μg LPS (*E. coli* strain O55:B5). Wet food was provided to animals in the days following LPS administration, due to expected diarrhea and weight loss. Animals reaching a scoring > 0.4 according to the Karolinska Institutet humane endpoint guidelines were removed from the experiment. Macroscopic signs of arthritis were assessed via a 0-60 scoring system based on the number of inflamed joints: 1 point per toe, 1 point per knuckle and 5 points for an inflamed wrist or ankle, totaling a maximum of 15 points per paw, or 60 points per animal. Patient-derived monoclonal ACPAs (1.5 mg per mouse) were co-transferred at day 0 together with the anti-CII antibody cocktail or at the peak of disease (day 7-8) to assess their inflammatory properties. Similarly in other experiments, CCP-pools from RA patients, the respective flow-through (FT) pool or a control IVIG were transferred to mice (2 mg per mouse) at the time of CAIA induction. All animal experiments were conducted under ethical permits approved by the local ethical committees (Stockholm, Sweden).

*Generation of F(ab')2 fragments*

Preparation of F(ab')2 fragments from the C03 monoclonal antibody was performed with a Pierce F(ab')2 preparation kit according to the manufacturer instructions (ThermoFisher Scientific). Briefly, a solution of C03 monoclonal antibody was initially desalted using a Zeba Spin Desalting Column. Digestion was achieved after 6h incubation of the monoclonal antibody with a freshly immobilized pepsin column. Undigested IgG and F(ab')2 were separated via a Nab protein A Plus Spin column, with confirmation of the purified F(ab')2 fragment by SDS-PAGE.

*Patient-derived CCP- and flow-through-pools*

The collection of patient-derived CCP-pool and their respective flow-through pools were prepared as previously described[8]. Briefly, plasma was obtained from peripheral blood samples of RA patients (n=37; ≈10 ml/patient) and centrifuged at 3000 g for 5 min. IgGs were purified on HiTrap Protein G HP columns (Cytiva), according to the manufacturer's instructions. After low pH elution and neutralization, IgG was dialyzed against PBS and applied to a CCP2 affinity column (kindly provided by Euro-Diagnostica, Sweden). ACPA were then obtained by elution of the column with 0.1 M glycine–HCl buffer (pH 2.7), and the pH was immediately adjusted to 7.4 with 1 M Tris (pH 9). The flow-through (FT) containing the unbound IgG fraction was also collected and equally dialyzed against PBS. Both ACPA and FT fractions were concentrated and buffer exchanged to PBS using 10 kDa spin Amicon centrifugation units (ThermoFisher Scientific) immediately prior to in vivo usage. ACPA was quality controlled by anti-CCP2 ELISA (Immunoscan CCPlus assay, Euro-Diagnostica, Sweden), SDS-PAGE/ Coomassie Brilliant Blue staining, size exclusion and LAL endotoxin testing.

*Flow cytometry analysis*

Mice undergoing CAIA were bled one drop of blood every 3rd day. After lysis of the red blood cells, the remaining cellular components were stained with a viability dye (Zombie NIR fixable dye; Biolegend) for 15min on ice. Cells were washed with PBS and resuspended in a Brilliant Stain buffer solution containing an antibody cocktail of fluorescently labelled monoclonal antibodies (Pacific Blue anti-FceR1a, clone Mar-1; BV510 anti-CD11b, clone M1/70; BV605 anti-CD64, clone X54-5/7.1; BV650 anti-Ly6G, clone 1A8; BV711 anti-CD16.2, clone 9E9; BV785 anti-Ly6C, clone HK1.4; FITC anti-CD32, clone S17012B; Spark Blue 550 anti-mouse CD19, clone 6D5; PE anti-CD169, clone 3D6.112; PE-Dazzle 594 anti-CD16, clone S17014E; PerCP anti-CD45, clone 30-F11; PE-Cy7 anti-CD11c, clone N418; APC anti-F4/80, clone BM8; Spark NIR 685 anti-CD117, clone 2B8; Alexa Fluor 700 anti-I-A/I-E, clone M5/114.15.2; all from Biolegend). After a 15min incubation on ice, cells were washed with PBS containing 0.5% bovine serum albumin and 2mM EDTA. Cells were fixed with eBioscience Foxp3/transcription factor fixation buffer according to the manufacturer instructions (ThermoFischer Scientific), acquired in a 3-laser spectral flow cytometer (Cytek Biosciences) and analyzed with FlowJo software (BD Biosciences).

*Reactivity against citrullinated peptides and proteins*

ACPA mAbs were screened for binding to citrullinated, native control peptides and control antigens using a custom-designed antigen microarray (ThermoFisher Scientific, Immunodiagnostics)[9]. ELISA reactivity to modified citrullinated vimentin peptides (Cit-Vim58-69 GRVYAT-Cit-SSAVR) were evaluated using plates from Orgentec Diagnostika, and citrullinated Histone 4 using biotinylated peptides (Cit3-His4 bio: SG[Cit]GKGGKGLGKGGAKRHGSGSK-biotin) on streptavidin-coated high-capacity pre-blocked plates (Thermo Fisher Scientific) and hIgG1 at 5 μg/ml in RIA buffer (1% BSA, 325 mM NaCl, 10 mM Tris-HCl, 1% Tween-20, 0·1% SDS) and detection with HRP conjugated Fab'2 goat anti-human IgG(γ) (Jackson Immunoresearch) and TMB substrate (Biolegend). Antibody binding to full-length citrullined protein was evaluate by on-plate modification and ELISA. Briefly, antigen were diluted to 3 μg/ml in 8M Urea and coated to high-bind half area plates (Corning). Plates were blocked with 5% low-fat milk in PBS and antigens citrullinated with 280mU/ml recombinant human PAD4 (Cayman chemicals) in 50 mM Tris 10 mM $CaCl_2$

1 mM DTT for 3h at 37C. Binding was assayed at 5 μg/ml hIgG1 to citrullinated vimentin, alpha enolase (in-house expressed human) and collagen II (bovine purified, chondrex) and at 0.5 μg/ml hIgG1 to citrullinated fibrinogen (human purifed, Sigma Aldrich).

**Supplemental Table 2 – Citrulline peptides analyzed on array**

| Peptide | Protein | Sequence* | Reference |
|---|---|---|---|
| Fibα 36-50 | Fibrinogen α-chain | GP(cit)VVF(cit)HQSACKDSK | [10,11] |
| Fibβ 36-52 | Fibrinogen β-chain | NEEGFFSA(cit)GHRPLDKK | [12] |
| Fibβ 60-74 | Fibrinogen β-chain | (cit)PAPPPISGGGY(cit)A(cit) | [10,11] |
| Fil 307-324 | Filaggrin | HQCHQEST(cit)GRSRGRCGRSGS[cyclic] | [13] |
| TNC1 2026-2040 | Tenascin | VFLRRKNG(cit)ENFYQNW | [14] |
| TNC5 2176-2200 | Tenascin | EHSSIQFAEMKL(cit)PSNF(cit)NLEG(cit)(cit)K(cit) | [14] |
| Vim 60-75 | Vimentin | VYAT(cit)SSAV(cit)L(cit)SSVP | [12] |

*The peptide amino acid sequence displayed where arginine R is substituted with citrulline in (cit). Both citrulline containing peptides and native arginine containing peptides were also analyzed on the array.
Besides native arginine peptides the following control autoantigens were included on the array: CENP B, collagen II, fibrillarin, Jo-1, Mi-2, PCNA, PM-Scl 100, Rip P0. Rip P1, Rip P2, RNA Pol III, RNP-70, RNP-A, RNP-C, Ro52, Ro60, Scl-70, SmBB, Sm, SSB/La
The ACPA fine-specificity array is further described in[9,15].

*CCP-ELISA*

The development of ACPA during the development of CAIA was tested through a CCP2-ELISA (Immunoscan CCPlus, Svar Life Science) with some modifications. Briefly, different serum dilutions from naïve mice or mice submitted to CAIA (day 15) were applied to a CCP pre-coated plate. Detection of murine anti-CCP-reactive antibodies was done with a peroxidase-conjugated F(ab')2 fragment goat anti-mouse IgG (Jackson ImmunoResearch) and using TMB as the enzymatic substrate. The arthritogenic cocktail of anti-CII antibodies used to induce CAIA was similarly tested in the same assay, in order to investigate any potential cross-reactivity of these antibodies with citrullinated antigens.

*C1q-ELISA*

The capacity of each ACPA clone in activating the classical pathway of complement was assessed by a C1q ELISA. A 96 well plate (Nunc, Maxisorp) was coated with each ACPA clone individually, starting at a concentration of 10μg/ml and followed by a stepwise two-fold dilution. Blocking was performed with a 2% ovalbumin solution. Purified mouse C1q protein (Complement Technologies Inc, USA) was then added to the wells at 1µg/ml followed by detected with a biotin-conjugated anti-mouse C1q antibody (ThermoFisher Scientific, clone JL-1). Streptavidin-HRP and TMB were used as enzyme and substrate of the reaction, respectively. The plate was washed 3-5 times between each step with PBS containing 0.05% Tween-20.

*Preparation and nanoLC-MS/MS analysis of mAb*

The purified mAb preparations (30 µg each, in 50 µl of 1% sodium deoxycholate (SDC), 50 mM triethylammonium bicarbonate (TEAB) pH 8.0) were reduced with 4.5 mM dithiothreitol (DTT) at 56 °C for 30 min and alkylated with 9 mM 2-iodoacetamide in the dark for 30 min at room temperature (RT). The alkylation reactions were quenched by incubation with DTT (9 mM final concentration) for 15 minutes at RT. Prior to proteolytic digestions, samples were diluted with 50mM TEAB to final volume of 90 µl. For each mAb preparation, 60 µl (20 µg) were incubated with trypsin (overnight at 37 °C, 0.2 µg, Pierce™ MS grade) and 30 µl (10 µg) with chymotrypsin (overnight at RT, 0.15 µg, Promega). SDC was removed by acidification with 10% trifluoroacetic acid (TFA) and subsequent centrifugation. The supernatants were further purified using Pierce peptide desalting spin columns (Thermo Fisher Scientific), according to the manufacturer's instructions. Purified preparations were reconstituted in 2% acetonitrile (ACN), 0.1% TFA for nanoLC-MS/MS analysis.

NanoLC-MS/MS was performed on an Orbitrap Exploris 480 mass spectrometer interfaced with Easy-nLC1200 liquid chromatography system (both Thermo Fisher Scientific). Peptides were trapped on an Acclaim Pepmap 100 C18 trap column (100 µm x 2 cm, particle size 5 µm, Thermo Fischer Scientific) and separated on an in-house packed analytical column (75 μm x 30 cm, particle size 3 μm, Reprosil-Pur C18, Dr. Maisch) using a gradient from 5% to 35% ACN in 0.2% formic acid over 75 min at a flow of 300 nL/min. Each preparation was analyzed using two different MS1 scans settings, m/*z* 380-1500 and *m/z* 600-2000, both at a resolution of 120K. MS2 analysis was performed in a data-dependent mode at a resolution of 30K, using a cycle time of 2 seconds. The most abundant precursors with charges 2–7 were selected for fragmentation using HCD at collision energy settings of 30. The isolation window was set to either 1.2 *m/z* for data acquired in *m/z* 380-1500 range or 3.0 *m/z* for data acquired in *m/z* 600-2000 range. The dynamic exclusion was set to 10 ppm for 20 s.

The acquired data were analyzed using Proteome Discoverer 2.4 (Thermo Fisher Scientific). Database searches were performed with Byonic (Protein Metrics) as search engine against custom protein database consisting of the selected mAb sequences. Precursor mass tolerance was set to 10 ppm and fragment mass tolerance to 30 ppm. Tryptic peptides with up to two missed cleavages and chymotryptic peptides with up to five missed cleavages were accepted. Custom glycan database consisted of 72 compositions, including sialylated and mono-fucosylated structures. Additional variable modification of methionine oxidation and fixed cysteine alkylation were allowed. Target decoy was used for peptide spectrum match (PSM) validation. Prior to the final assignment, the identified glycosylated peptides were manually evaluated based on the observed fragmentation pattern, the number of glycoforms per site, the number of PSM per glycoform, and the retention time windows for the different glycoforms (at the same site). The extracted ion chromatogram peak intensities were used to determine the glycoform abundances and are expressed as percent of total signal for all modified and non-modified peptides sharing the same amino acid sequence.

*Experimental testing and statistical analysis*

All animal experimental settings were repeated at least twice, with a total number of 10 to 50 animals assessed per condition, depending on the ACPA clone used. A minimal of 5 animals per group was used at any given time given the prevalence of arthritis being generally 80-100%, thus allowing statistical calculations with a good degree of power, and assuming a non-normal distribution of the data. Each cage contained no more than two animals of the same experimental group, with all experimental groups assigned to as many cages in order to minimize potential cage effects. Statistical analysis of arthritis development and prevalence between groups was conducted by non-parametric repeated-measures Friedman test with Dunn's multiple comparison test. When comparing two or more groups in single observation assays, a non-parametric Mann-Whitney or Kruskal-Wallis test was used. Prism GraphPad version 9.3.1 was used to calculate statistics and statistical significance was considered when $p < 0{\cdot}05$ for a 95% confidence interval.

References

1. Amara, K. *et al.* A Refined Protocol for Identifying Citrulline-specific Monoclonal Antibodies from Single Human B Cells from Rheumatoid Arthritis Patient Material. *Bio-protocol* **9**, e3347 (2019).

2. Sahlström, P. *et al.* Different Hierarchies of Anti–Modified Protein Autoantibody Reactivities in Rheumatoid Arthritis. *Arthritis Rheumatol* **72**, 1643–1657 (2020).

3. Steen, J. *et al.* Recognition of Amino Acid Motifs, Rather Than Specific Proteins, by Human Plasma Cell–Derived Monoclonal Antibodies to Posttranslationally Modified Proteins in Rheumatoid Arthritis. *Arthritis Rheumatol* **71**, 196–209 (2019).

4. Lloyd, K. A. *et al.* Variable domain N-linked glycosylation and negative surface charge are key features of monoclonal ACPA: Implications for B-cell selection. *Eur J Immunol* **48**, 1030–1045 (2018).

5. Titcombe, P. J. *et al.* Pathogenic Citrulline-Multispecific B Cell Receptor Clades in Rheumatoid Arthritis. *Arthritis Rheumatol* **70**, 1933–1945 (2018).

6. Bournazos, S. *et al.* Broadly Neutralizing Anti-HIV-1 Antibodies Require Fc Effector Functions for In Vivo Activity. *Cell* **158**, 1243–1253 (2014).

7. Unger, T., Jacobovitch, Y., Dantes, A., Bernheim, R. & Peleg, Y. Applications of the Restriction Free (RF) cloning procedure for molecular manipulations and protein expression. *J Struct Biol* **172**, 34–44 (2010).

8. Ossipova, E. *et al.* Affinity purified anti-citrullinated protein/peptide antibodies target antigens expressed in the rheumatoid joint. *Arthritis Res Ther* **16**, R167 (2014).

9. Hansson, M. *et al.* Validation of a multiplex chip-based assay for the detection of autoantibodies against citrullinated peptides. *Arthritis Res Ther* **14**, R201 (2012).

10. Iobagiu, C. *et al.* The antigen specificity of the rheumatoid arthritis-associated ACPA directed to citrullinated fibrin is very closely restricted. *J Autoimmun* **37**, 263–272 (2011).

11. Sebbag, M. *et al.* Epitopes of human fibrin recognized by the rheumatoid arthritis-specific autoantibodies to citrullinated proteins. *Eur J Immunol* **36**, 2250–2263 (2006).

12. Verpoort, K. N. *et al.* Fine specificity of the anti–citrullinated protein antibody response is influenced by the shared epitope alleles. *Arthritis Rheumatism* **56**, 3949–3952 (2007).

13. Schellekens, G. A. *et al.* The diagnostic properties of rheumatoid arthritis antibodies recognizing a cyclic citrullinated peptide. *Arthritis Rheumatism* **43**, 155–163 (2000).

14. Schwenzer, A. *et al.* Identification of an immunodominant peptide from citrullinated tenascin-C as a major target for autoantibodies in rheumatoid arthritis. *Ann Rheum Dis* **75**, 1876 (2016).

15. Rönnelid, J. *et al.* Anticitrullinated protein/peptide antibody multiplexing defines an extended group of ACPA-positive rheumatoid arthritis patients with distinct genetic and environmental determinants. *Ann Rheum Dis* **77**, 203 (2018).
