## Supplementary Figures for "Divergent and dominant anti-inflammatory effects of patient-derived anti-citrullinated protein antibodies (ACPA) in arthritis development"

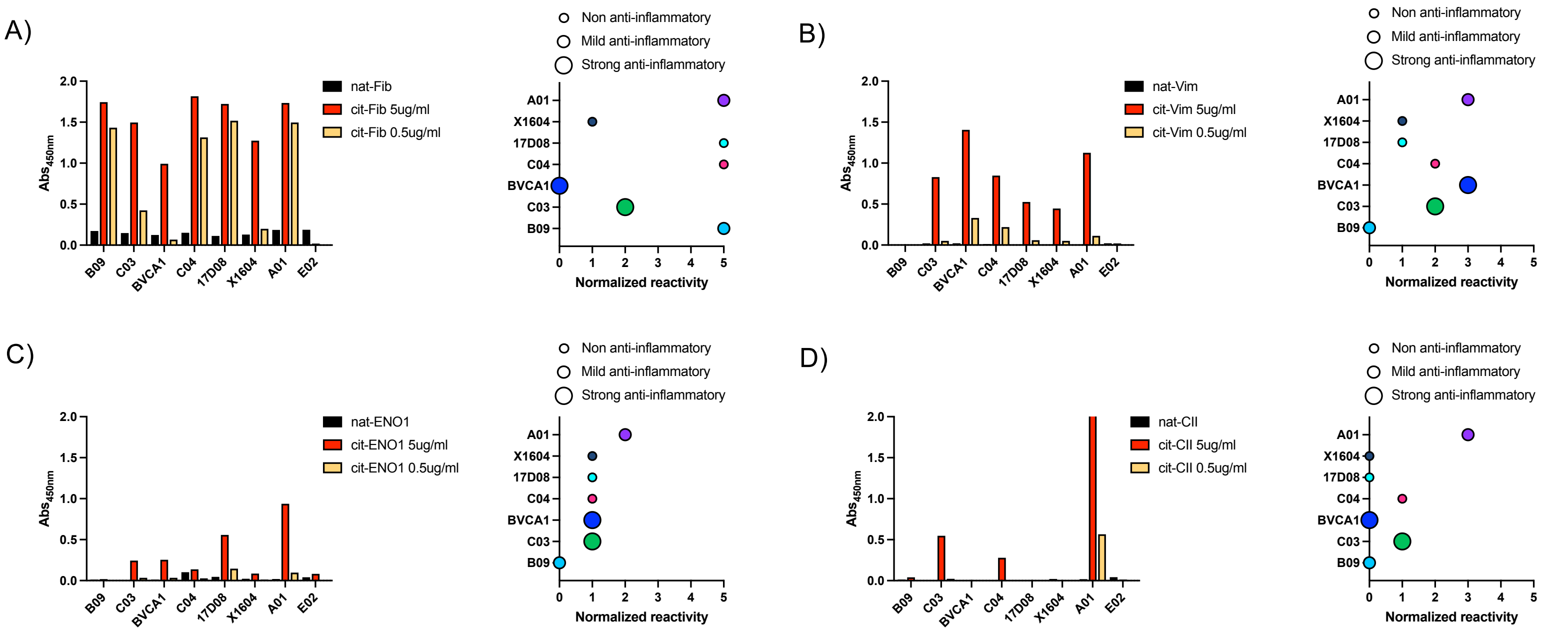


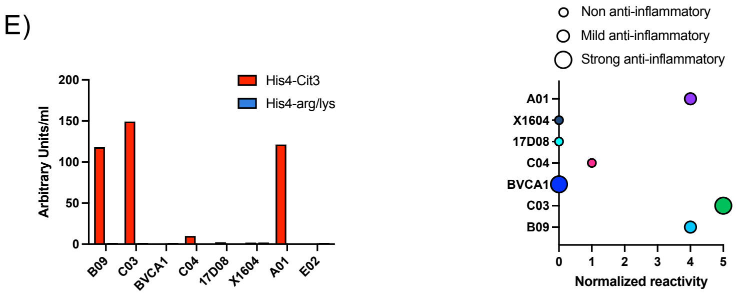


**Supplementary figure 1 – Citrullinated antigen reactive fine-specificities by monoclonal ACPA do not associate with observed anti-inflammatory effects.** Reactivity to citrullinated and native full protein fibrinogen (A), vimentin (B), alpha-enolase (C) and type II collagen (D) by monoclonal ACPAs and respective associations with observed anti-inflammatory effects in CAIA. (E) Reactivity of tested ACPA clones against the citrulline 3 peptide from histone 4 (His4) identified by Chirivi and colleagues [*R. Chirivi et al (2021); Therapeutic ACPA inhibits NET formation: a potential therapy for neutrophil-mediated inflammatory diseases; Cellular and Molecular Immunology 18:1528*]. Nat-Fib: native fibrinogen; cit-Fib: citrullinated fibrinogen; nat-Vim: native vimentin; cit-Vim: citrullinated vimentin; nat-ENO1: native alpha-enolase; cit-ENO1: citrullinated alpha-enolase; nat-CII: native type II collagen; cit-CII: citrullinated type II collagen.

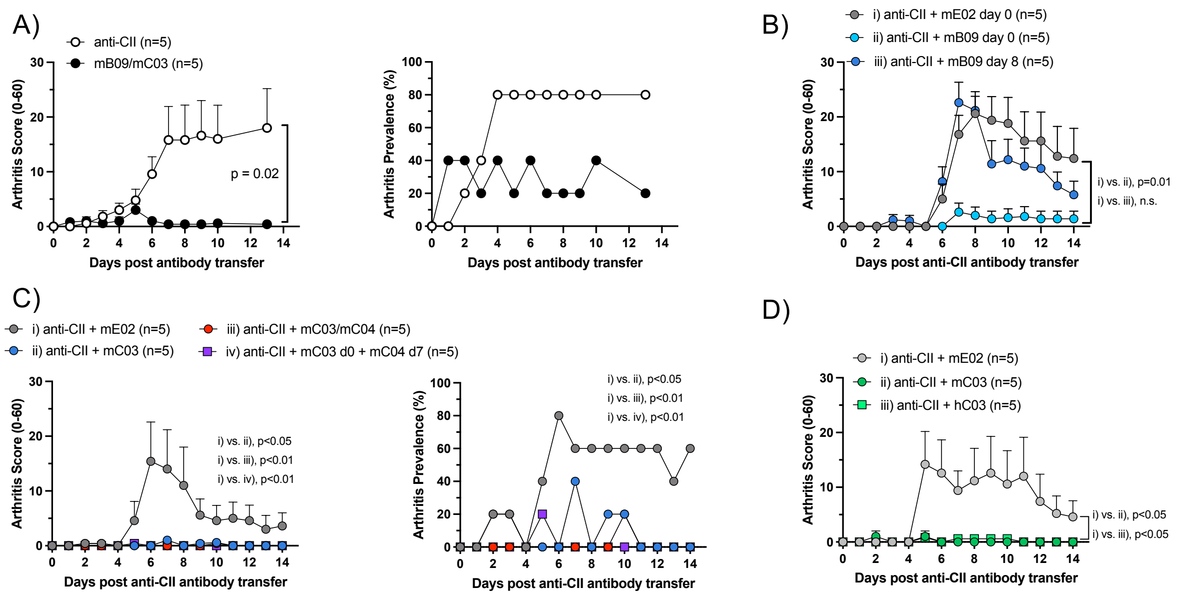


**Supplementary Figure 2 – Supporting observations on how ACPAs affect inflammatory arthritis in vivo.** (A) Transfer of anti-CII antibody cocktail or a combination of two monoclonal ACPAs i.v. day 0, followed by LPS administration i.p. day 4. (B-D) CAIA was induced in mice by i.v. transfer of arthritogenic anti-CII antibody cocktail. Boosting and synchronization of disease symptoms was done by i.p. administration of LPS 3 days post disease induction. (B) mB09 ACPA transferred at day 0 or 7 to assess therapeutic potential. (C) Co-administration of anti-inflammatory mC03 and disease-prone mC04 ACPAs at day 0, or mC03 day 0 followed by mC04 day 7. (D) Comparison between C03 ACPA expressing a murine IgG2a or human IgG1 constant domain. Statistical analysis calculated by non-parametric repeated-measures Friedman test with Dunn's multiple comparison test.

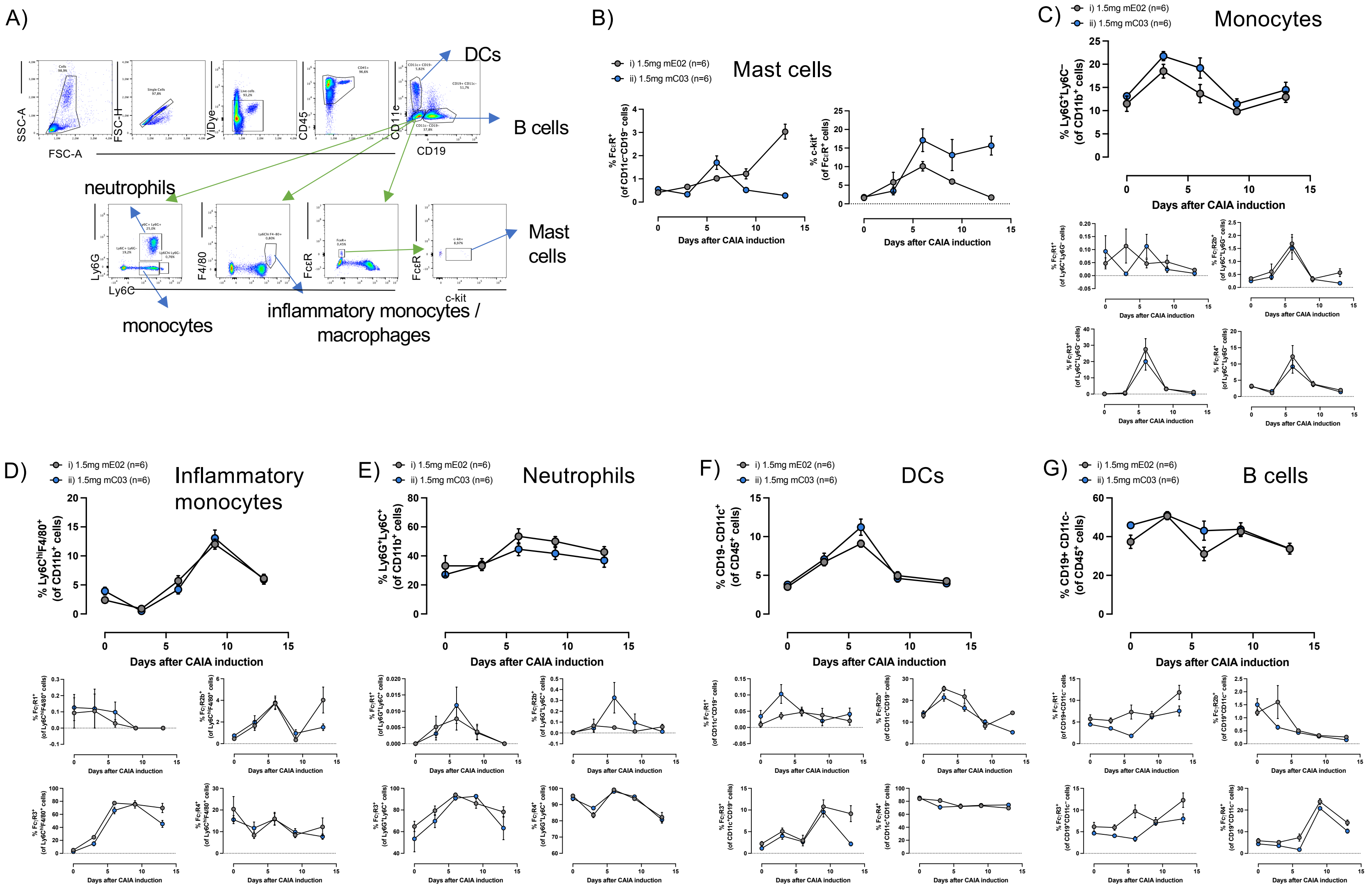


**Supplementary figure 3 – Frequency of circulating immune cells and their expression of FcγR during CAIA is not affected by administration of ACPA.** CAIA was induced in BALB/c mice followed by LPS administration day 3. mE02 or mC03 ACPA were transferred at day 0. Blood was collected every 3rd day for evaluation of immune cell frequencies in the blood, as well as their corresponding expression of FcγR1, FcγR2b, FcγR3, FcγR4. (A) Gating strategy. (B) Representation of FcγR expression in mast cells (FcεR+ c-kit+), (C) monocytes (Ly6C+Ly6C–),(D) inflammatory monocytes/macrophages (Ly6Ghi F4/80+), (E) neutrophils (Ly6G+Ly6C+), (F) DCs (CD11c+ CD19–) and (G) B cells (CD19+ CD11c–). No differences in mean fluorescence intensity (MFI) of any of the FcγR were detected between mice receiving mE02 or mC03.

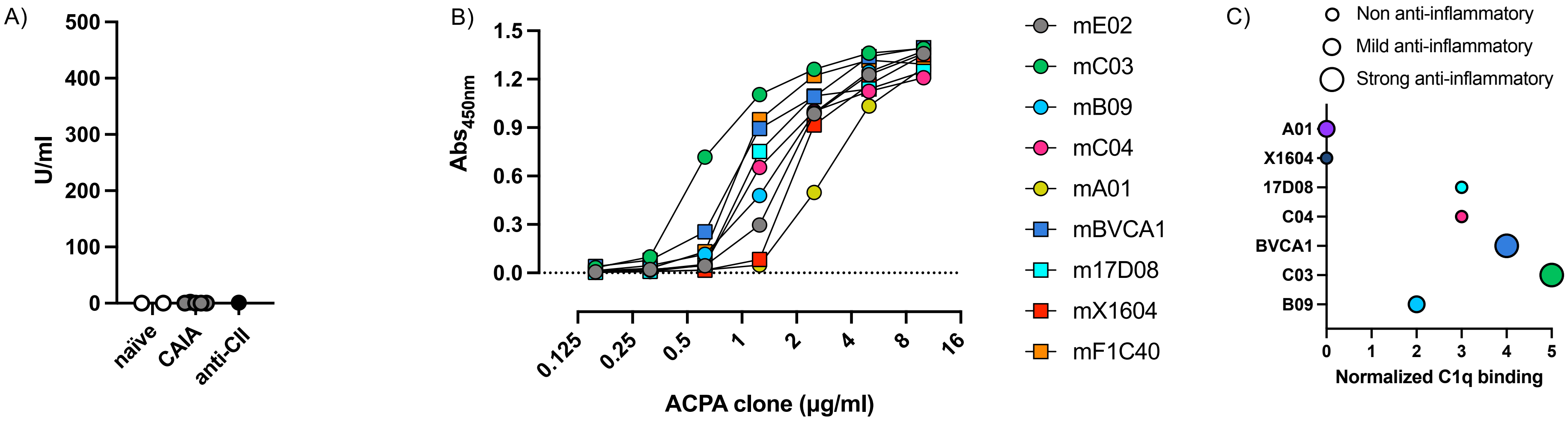


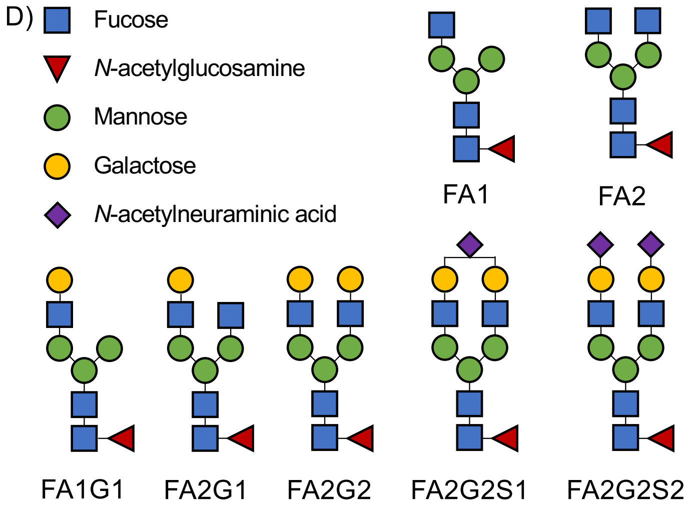


E)

| ACPA | FA1 | FA2 | FA1G1 | FA2G1 | FA2G2 |
|---|---|---|---|---|---|
| mE02 | n.d. | 56 | n.d. | 15 | 1 |
| mB09 | 18 | 69 | n.d. | 12 | n.d. |
| mC03 | 8 | 58 | 2 | 30 | n.d. |
| mC04 | 15 | 59 | 2 | 13 | n.d. |
| mBVCA1 | 21 | 53 | 2 | 15 | n.d. |
| m17D08 | 14 | 50 | 10 | 10 | 1 |
| mX1604 | 20 | 54 | 4 | 12 | 2 |
| mF1C40 | 17 | 51 | 4 | 16 | 2 |
| mA01 | 18 | 51 | 2 | 17 | 3 |

**Supplementary figure 4 – Glycosylation forms and complement deposition by ACPA clones do not clearly explain observed anti-inflammatory effects.** (A) Anti-citrulline reactivity was determined through a CCP-ELISA in serum from naïve mice, mice that were submitted to CAIA (day 15 post disease induction), as well as the anti-CII arthritogenic antibody cocktail used to induce the arthritis model. (B) Complement C1q deposition induced by different ACPA clones. (C) Representation of the normalized C1q binding shown in (B) at a concentration of 1.25µg/ml monoclonal ACPA. (D) Schematic representation of major Fc glycoforms. FA2G2S1 and FA2G2S2 glycoforms were not found in any of the ACPA clones. (E) Relative abundance of each glycoform found in the ACPA clones used in vivo.
